# T cell stiffness is enhanced upon formation of immunological synapse

**DOI:** 10.1101/2021.01.06.425650

**Authors:** Philipp Jung, Xiangda Zhou, Markus Bischoff, Bin Qu

**Affiliations:** Institute for Medical Microbiology and Hygiene, Saarland University, Homburg, Germany; Department of Biophysics, Center for Integrative Physiology and Molecular Medicine (CIPMM), School of Medicine, Saarland University, Homburg; Leibniz Institute for New Materials, Saarbrücken, Germany

## Abstract

T cells are activated by cognate target cells via an intimate contact, termed immunological synapse (IS). Cellular mechanical properties, especially stiffness, are essential to regulate cell functions, T cell stiffness at a subcellular level at the IS still remains largely elusive. In this work, we established an atomic force microscopy (AFM)-based elasticity mapping method on whole T cells to obtain an overview of the stiffness with a resolution of ~ 60 nm. Using Jurkat T-cells and primary human CD4^+^ T cells, we show that in the T cells in contact with functionalized surfaces, the lamellipodia are stiffer than the cell body. Upon IS formation, T cell stiffness is substantially enhanced both at the lamellipodia and in cell body. Chelation of intracellular Ca^2+^ abolishes IS-induced stiffening at the lamellipodia but has no influence on cell body-stiffening, suggesting different regulatory mechanism of IS-induced stiffening between the lamellipodia and the cell body.

## Introduction

T cells belong to the adaptive immune system and can be classified into CD4^+^ T cells and CD8^+^ T cells. CD4^+^ T cells are essential for orchestrating immune responses and CD8^+^ T cells are the key players to eliminate tumor and pathogen-infected cells. T cells are activated by the engagement of T-cell receptors (TCR) with the matching antigen. Consequently, CD3 molecules, one key component of the TCR complex, transduce the signal to activate downstream pathways leading to formation of a tight junction between T cells and target cells termed the immunological synapse (IS). Upon IS formation, a fundamental rearrangement of cytoskeleton takes place, for example reorientation of microtubule-organizing center (MTOC) towards the IS and formation of F-actin ring structure at the IS. In between, the adhesion molecule LFA-1 (lymphocyte function-associated antigen 1) binds its ligand on the target cells to seal and stabilize the IS. Upon T cell activation, intracellular Ca^2+^ concentration is drastically enhanced via Ca^2+^ influx. Ca^2+^ serves as an essential second messenger in T cells to regulate their key functions such as activation, proliferation, and effector functions.

Mechanical properties, including stiffness and mechanical forces, play a significant role in modulation of cell functions, especially T cells (Harrison, Fang et al., 2019). It is reported that cytotoxic T lymphocytes optimize their killing function via applying mechanical forces (Basu, Whitlock et al., 2016). Later studies using micropillar arrays show that T cells exert lateral forces at the interface, where the mechanic output is coordinated with release of cytotoxic granules (Jin, Tamzalit et al., 2019, Tamzalit, Wang et al., 2019). In addition, the mechanical forces generated by T cells also include pushing and pulling forces in the piconewton (pN) range upon activation (Basu et al., 2016, Husson, Chemin et al., 2011). Force generation of T cells requires a sustained elevation of intracellular calcium, integrity of functional F-actin network including actomyosin contractility, and PI3K signaling (Basu et al., 2016, Hui, Balagopalan et al., 2015, Jin et al., 2019). Interestingly, even when actin-cytoskeleton is perturbed by latrunculin-A, application of periodical mechanical forces can still induced Ca^2+^ influx, when the forces are linked with TCR (Basu et al., 2016). Furthermore, T cells can sense stiffness of target cells or substrates and translate the mechano-signal to functional consequences. For example, responsiveness of T cells to stimuli is elevated on stiffer substrates (40-50 kPa) relative to their softer counterparts (< 12 kPa) (Majedi, Hasani-Sadrabadi et al., 2020, Zhang, Zhao et al., 2020 {Majedi, 2020 #35). How the stiffness of T cells *per se* is regulated upon IS formation is still not well understood.

During the last 20 years, many different approaches had been made to address the stiffness and stiffening processes of eukaryotic cells, especially immune cells. Early approaches like micropipette aspiration and microplate-based rheometry involved complex technical structures in which the cells were inserted or incorporated and mechanically stressed (Desprat, Guiroy et al., 2006, Hochmuth, 2000). Cytometry-related approaches are basing on principles like deformation during deceleration in flow or twisting triggered by membrane-bound metal beads in a magnetic field. Those methods are limited to a constant fluid stream to operate. In contrast, methods which are able to characterize elastic properties of cells in an environment modeling their natural habitat, like cell monolayers or mimicking cell-cell interactions, are capable to provide much deeper insights into biomechanical processes. Optical tweezers use focused laser beams to generate attractive or repulsive forces in the pN range, and are considered promising to characterize and modify biomaterials. Hence, this method is mostly used to move or sort cells and is still not broadly applied in characterizing local cell properties (Killian, Ye et al., 2018, Wang, Butler et al., 1993). Atomic Force Microscopy (AFM) applies a mechanical probe, often made of silicon or silicon nitride with a nanometer-thin tip. This probe is moved along the cell surface and is interacting with each point of a defined scanning grid. Hence, it allows the examination of height profiles or measure mechanical properties with nanometer spatial and pN force resolution (Thewes, Loskill et al., 2015). In recent years, AFM has evolved as one of the most important tools in cell biology to study adhesive forces, cell deformation, cell turgor, and cell elasticity properties (Loskill, Pereira et al., 2014, Pi & Cai, 2019, Scheuring & Dufrene, 2010).

In this work, we established an AFM-based method to simultaneously characterize surface profile and stiffness of live CD4^+^ T cells. T cells were attached to glass coverslips via adhesion molecule LFA-1 with or without CD3/CD28 activation. We found that lamellipodia were dynamically formed upon T cells got in contact with the antibody-coated coverslips independent of CD3/CD28 activation. At the lamellipodia, the stiffness is significantly higher than that at the cell body. Remarkably, upon formation of IS induced by CD3/CD28 stimulation, T cells were substantially stiffened at the cell body as well as at the lamellipodia. Calcium is involved in regulation of this IS formation-induced T cell stiffening. Our results show that the stiffness of T cells is significantly increased upon IS formation, which could facilitate T cells to generate and exert mechanical force on their target cells.

## Results

### T cells are stiffened upon IS formation

To examine the stiffness of T cells upon formation of the IS, we established a method to investigate live T cells on functionalized coverslips by AFM based Peak Force Quantitative Nanoscale Mechanical Characterization (Peak Force QNM) (Berquand, C. Roduit et al., 2010). We first functionalized polyornithine-coated coverslips with anti-CD3- and anti-CD28 antibodies, which activate the TCR of T cells and related downstream pathways via CD3 and the co-stimulation molecule CD28 (Qu, Pattu et al., 2011). Jurkat T cells, which present features of effector CD4^+^ T cells (Abraham & Weiss, 2004), were next settled on the functionalized coverslip and allowed to interact with the antibodies for 15 min. Subsequently, a quarter of the each cell was investigated by Peak Force QNM at a resolution of ~60 nm between measurements to create a high density map of local elastic moduli (Youngs moduli determined by Derjaguin-Muller-Toporov fit). We observed that Jurkat T cells formed an IS with the antibodies deposited on the coverslip, by forming clear lamellipodial protrusions, which dynamically changed over time (Figs. 1A, B). This morphology of lamellipodia detected by AFM resembles what have been observed from scanning electron microscopy and immunostaining (Saitakis, Dogniaux et al., 2017, Schoppmeyer, Zhao et al., 2017). In addition, Complete height profiles were also obtained, showing that T cells were flattened after contacting with the functionalized surface (Fig. 1C), suggesting a functional IS was formed (Pattu, Qu et al., 2011, Qu et al., 2011).

**Figure 1.**
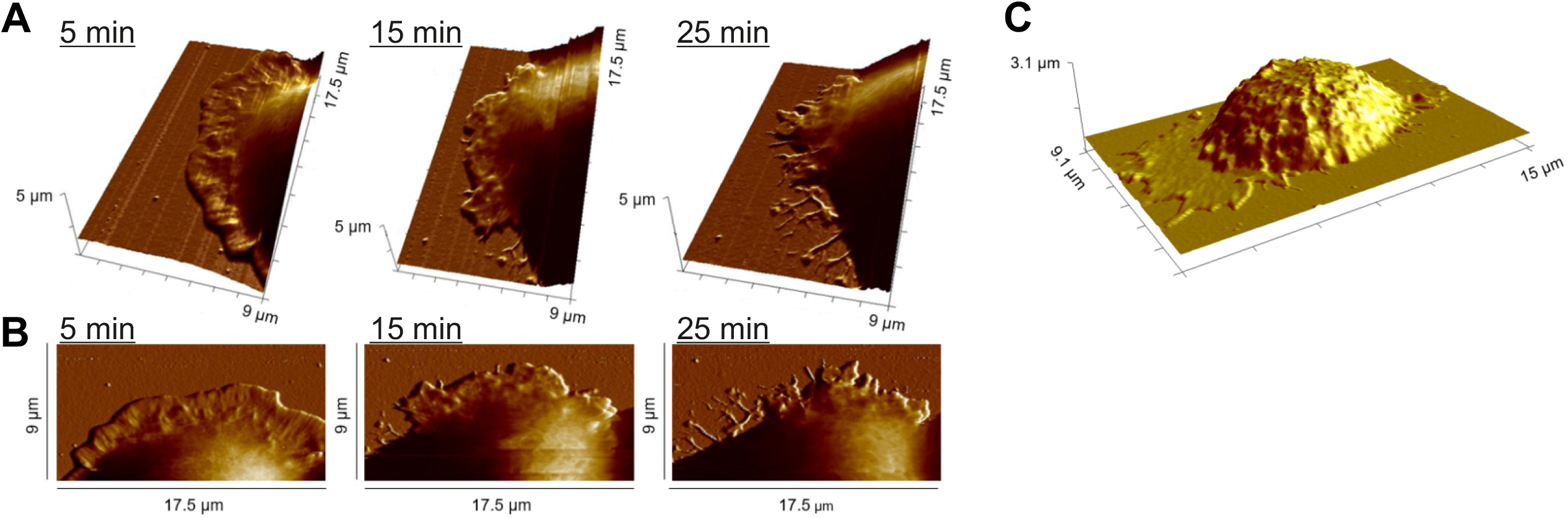
Dynamics of lamellipodium formation at the IS; **A, B:** Dynamic changes of the lamellipodium of a Jurkat T cell during IS formation on a αLFA-1+αCD3+αCD28 antibody-coated coverslip. The height profile, examined by Peak Force QNM, is displayed (A: 3D view, B: top view). **C:** Height profile of a whole Primary T cell during IS formation on an antibody-coated coverslip.

Next, to investigate the impact of IS formation on T cell stiffness, we placed Jurkat T cells on either anti-LFA-1 antibody-decorated surfaces or anti CD3-/CD28-/LFA-1 antibodies-coated surfaces. Here we involved anti-LFA-1 antibody for three reasons. Firstly, for the control condition, without anti-LFA-1 antibody T cells did not attach to the surface, which is a prerequisite to determine cell stiffness. Secondly, in physiological scenarios, LFA-1 is always engaged with its interaction partner ICAM-1 along with TCR activation. Thirdly, it is reported that spreading of T cells upon mechanosensing is significantly different when CD3 is activated alone or in combination with LFA-1 engagenment (Wahl, Dinet et al., 2019). Our results show that when Jurkat T-cells were brought into contact with the anti CD3-/CD28-/LFA-1 antibodies-decorated surfaces, stiffness of the cells was substantially enhanced at both the cell body and the lamellipodial protrusions (factor 11 and factor 23, respectively), when compared to Jurkat T-cells seeded on the LFA-1 antibody functionalized control surface (Figs. 2A-D). Jurkat cells also displayed on both substrates a significantly higher stiffness of the lamellipodial regions when compared to the cell body (Figs. 2E, F), which is in line with earlier observations made with different eukaryotic cell types (Innocenti, 2018).

**Figure 2.**
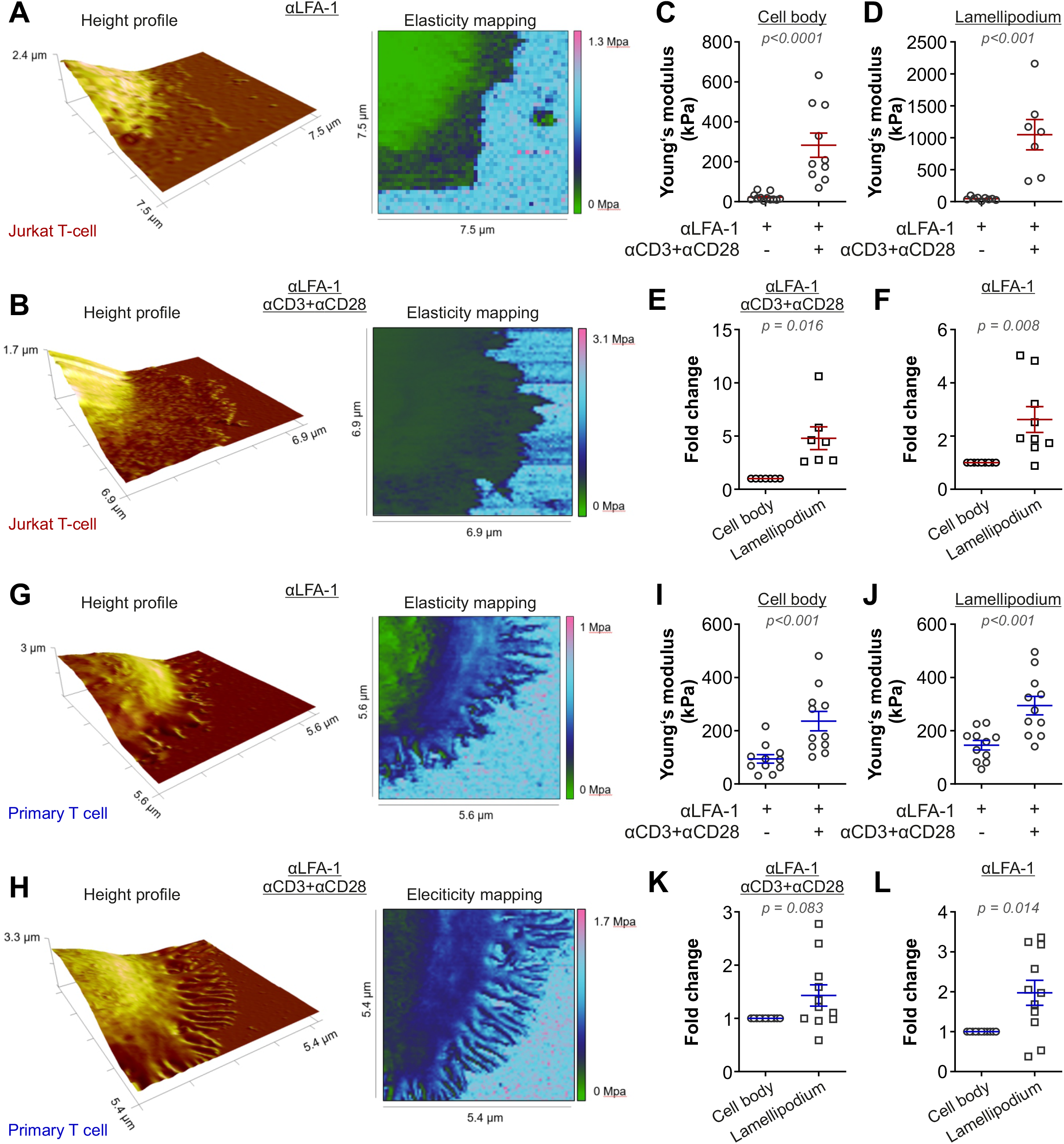
The stiffness of T cells increases upon activation; **A, B:** Height profiles and according elasticity maps of Jurkat T cell sections on αLFA-1 and αLFA-1+αCD3+αCD28 antibody-coated coverslips. **C, D:** Young’s moduli of cell bodies and lamellipodia of Jurkat T cells after IS formation on both antibody substrates described above. The Young’s moduli were obtained from 8 individual cells (αLFA-1) and 7 individual cells (αLFA-1+αCD3+αCD28), respectively. **E, F:** Fold changes between Young’s moduli of cell bodies and lamellipodia of Jurkat T cells after IS formation on both antibody substrates described above. **G, H:** Height profiles and according elasticity maps of primary human CD4^+^ T cell sections on αLFA-1 and αLFA-1+αCD3+αCD28 antibody-coated coverslips. **I, J:** Young’s moduli of cell bodies and lamellipodia of primary human CD4^+^ T cells after IS formation on both antibody substrates described above. The Young’s moduli were obtained from 10 individual cells (αLFA-1) and 11 individual cells (αLFA-1+αCD3+αCD28), respectively. **K, L:** Fold changes between Young’s moduli of cell bodies and lamellipodia of primary T cells after IS formation on both antibody substrates described above. The Wilcoxon test was used to test for statistically significant differences; individual p values are given in the plots. Results are from 11 independent experiments.

To exclude that the IS mediated changes in stiffness were not specific to Jurkat T cells, an additional series of measurements with unstimulated primary human CD4^+^ T cells was carried out (Figs. 2G-L). Primary human CD4^+^ T cells attaching to anti CD3-/CD28-/LFA-1 antibodies-decorated surfaces also exhibited a significantly enhanced stiffness at the lammelipodial regions and cell body when compared to comparable regions of primary human CD4^+^ T cells that attached to a LFA-1 functionalized surface (Figs. 2I, J), although fold changes in stiffness were much smaller (2,5-fold and 2-fold, respectively) than those seen with Jurkat T cells (Figs. 2E, F). The stiffness of the lamellipodial regions of primary human CD4^+^ T cells was on average again higher than that of the cell body (Figs. 2K, L), however, this difference was not statistically significant when a functional IS has been formed (Fig. 2K). Taken together, our findings prove that T cells are stiffened upon IS formation.

During analysis, we noticed that lamellipodia formed by T cells exhibited two morphologies, an extended and flattened form (e.g. Fig. 2A, B) and a needle-shaped filopodia-like form (e.g. Fig. 2G, H). Among Jurkat cells the extended morphology is more common (full antibody: 6 out of 7 cells; LFA-1: 8 out of 9 cells). Among primary CD4^+^ T cells, filopodia-like form is more frequent (full antibody: 6 out of 11 cells; LFA-1: 6 out of 11 cells). Interestingly, in the primary T cells, the filopodia on the substrates with full antibodies were significantly stiffer compared to those on the substrates with only LFA-1 antibody (p = 0.011). No statistical differences was found between flattened and filopodia forms.

We then asked whether the stiffness of lamellipodia measured by this method could be influenced by stiffness of the glass coverslip. During elasticity mapping a peak force threshold of 700 pN was applied which caused an indentation of 21 nm (± 2.2) into the cell body and 17 nm (± 4.7) into the lamellipodia, respectively. Since the lamellipodia revealed a mean height of 144 nm (± 69.7; with a maxium height of 335 and a minimum height of 51 nm) we can exclude an impact of the glass cover slip material on the Young’s moduli examined for the lamellipodia.

### T cell stiffening is regulated by calcium

We next sought for the underlying mechanism regulating T cell stiffening triggered by TCR-activation. Since the cytoskeleton plays an important role in maintaining cell stiffness (Gavara & Chadwick, 2016), we first targeted the actin-cytoskeleton as well as the microtubule-network by adding latrunculin-A (which inhibits actin polymerization and accelerates actin filament depolimeization) and nocodazole (a microtubule depolymerizing agent), respectively. Not surprisingly, with the disassembly of the cytoskeleton, primary human CD4^+^ T cells failed to form an IS on anti CD3-/CD28-/LFA-1 antibodies-decorated surfaces, and were also unable to firmly attach to the coverslip, which made it impossible to measure the corresponding stiffness by the methodology used here. Next, we turned our focus to calcium (Ca^2+^) by chelating intracellular Ca^2+^ with EGTA-AM, since TCR activation elicits a Ca^2+^ influx into T cells. We found that the lamellipodia formed by primary human CD4^+^ T cells on anti CD3-/CD28-/LFA-1 antibodies-decorated surfaces in presence of EGTA exhibited substantially lower stiffness, when compared to their DMSO-treated counterparts (Figs. 3A-D). Notably, the stiffness of the cell body was not affected by Ca^2+^ chelation (Fig. 3C), indicating that Ca^2+^ is not involved in regulating stiffening of the cell body upon IS formation.

**Figure 3.**
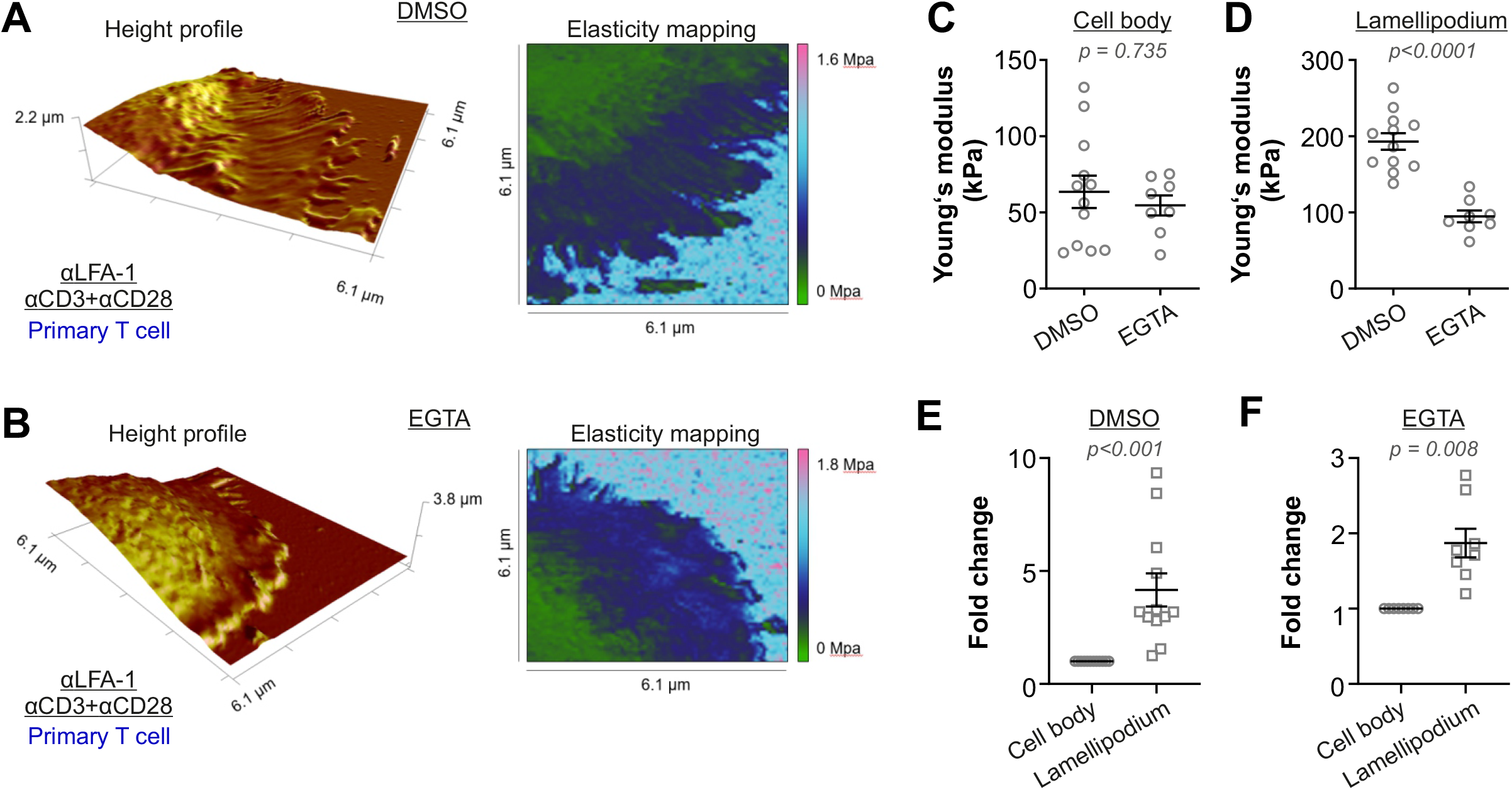
Activation-induced T cell stiffening is regulated by intracellular calcium. **A, B:** Height profiles and according elasticity maps of primary T cell sections on αLFA-1+αCD3+αCD28 antibody-coated coverslips. The cells were treated with EGTA-AM (dissolved in DMSO) to chelate intracellular Ca2^+^ (B) or treated with DMSO as a control (A). **C, D:** Young’s moduli of cell bodies and lamellipodia of primary T cells after IS formation on a αLFA-1+αCD3+αCD28 antibody-coated coverslip, treated with EGTA-AM or DMSO (control). The Young’s moduli were obtained from eight (EGTA-AM) and eleven individual cells (DMSO), respectively. **E, F:** Fold changes between Young’s moduli of cell bodies and lamellipodia of primary T cells after under the conditions described above. The Wilcoxon test was used to test for statistically significant differences; individual p values are given in the plots. Results are from 4 independent experiments.

## Discussion

In cell biology the development of imaging techniques provided and is still providing tremendously valuable information about cell morphology as well as changes and adaptations in cell shape due to external stimuli. Optical microscopy is capable to provide images of living cells fastly and without much sample preparation. Yet, the resolution is limited by the wavelength of light. Scanning electron microscopy (SEM) is capable to reach resolutions in the nanometer range but it still requires intensive sample preparation or sample surface modifications. In contrast, AFM-based techniques are able to reach resolutions in the nanometer range on living cells in media, located in cell clusters, monolayers or tissues. Combined with the possibility to examine interactions happening between the cell surface and the silicon probe, like cell stiffness or adhesion, AFM offers unique possibilities to study cell mechanics. In this work we tried to play out these advantages and created detailed elasticity maps of T cells upon IS formation. One of the major innovations of our study is that we are able to distinguish between the stiffness of the cell body and those of the lamellipodium. To our knowledge this is the first work addressing different cell parts in detail during the processes of the T cell activation. However, other AFM-based methods had been used before to study the IS formation. Several studies have utilized Single cell force spectroscopy (attaching a whole T cell to an AFM cantilever) to examine the adhesion forces to antigen presenting cells (Hoffmann, Hosseini et al., 2011, Hosseini, Louban et al., 2009, Lim & Ricciardi-Castagnoli, 2012). A recent study addressed the impact of cortical actin on surface rigidity of adherent and non-adherent T cells (Razvag, Neve-Oz et al., 2019). Furthermore, Saitakis and colleagues investigated the impact of substrate stiffness on TCR activation and T cell migration (Saitakis et al., 2017). In addition, influence from the stiffness of the cell nucleus can be excluded based on investigate the parameters applied in this study.

In this work, we also postulated that perturbation of cytoskeleton would significantly influence T cell stiffening induced by IS formation both at the lamellipodia and the cell body. However, we failed to determine the role of cytoskeleton on T cell stiffening as disruption of the actin-cytoskeleton or of the microtubule network did not allow T cells to attach to the functionalized surface firmly enough for the cantilever to complete the scanning. Evidence shows that T cells with disassembled actin-cytoskeleton via latrunculin-A fail to generate mechanical forces (Basu et al., 2016). Fast growth of microtubules in CD4^+^ T cells at the contact site between anti-CD3 antibody-coated coverslip is observed, and the traction stresses at the IS generated by actomyosin contractility is increased after disassembly of the microtubule-network (Hui & Upadhyaya, 2017). Although little direct evidence for the relationship between cell stiffness of immune cells and cytoskeleton, studies on other cell types are inspiring. It is reported that for the stiffness of mouse cochlea hair cells and murine fibroblasts, actin-cytoskeleton is essential, whereas for the stiffness of supporting cells in mouse cochlea, microtubule-acetylation is decisive (Gavara & Chadwick, 2016, Szarama, Gavara et al., 2012). All these findings indicate that both actin- and microtubule-cytoskeleton could play a significant role in T cell stiffening upon activation. To confirm this, further experiments and new approaches must be carried out. Why is Ca^2+^ involved in regulating the stiffness at the lamellipodia but not at the cell body? Here we used EGTA-AM to chelate the intracellular Ca^2+^. When extracellular Ca^2+^ is chelated by EGTA, TCR activation can still induce Ca^2+^ release from the endoplasmic reticulum store, leading to a transient and moderate elevation of intracellular Ca^2+^, which is suffice to initiate some downstream events including actin polymerization and actin-dependent spreading (Babich & Burkhardt, 2013). However, the release of cytotoxic granules is substantially inhibited with very low extracellular Ca^2+^ (3 μM) (Zhou, Friedmann et al., 2018). Our result that chelation of intracellular Ca^2+^ reduces the stiffness only at the lamellipodia but not the cell body suggests that sustained or higher levels of intercellular Ca^2+^ are required to stiffen lamellipodia, and the stiffened lamellipodia could be important for vesicle fusion at the IS.

## Materials and Methods

### Antibodies and reagents

All chemicals not specifically mentioned are from Sigma-Aldrich (highest grade). The following antibodies or reagents were used: anti-Integrin alpha-L (ITGAL) antibody (Antibodies-online), mouse anti-human CD28 antibody (BD Pharmingen), and mouse antihuman CD3 antibody (Diaclone).

### Cell culture

Jurkat T-cells, Isolation and culture conditions of primary T lymphocytes:

Human peripheral blood mononuclear cells (PBMCs) were obtained from healthy donors as described before {Kummerow, 2014 #47}. Primary human CD4^+^ T cells were negatively isolated from the PBMCs using CD4^+^ T Cell Isolation Kit human (Miltenyl). The isolated CD4^+^ T cells were cultured in AIM V medium (ThermoFisher Scientific) with 10% FCS (ThermoFisher Scientific). Jurkat T-cells were cultured in RPMI-1640 medium (ThermoFisher Scientific) with 10% FCS. All cells were cultured at 37°C with 5% CO_2_.

### Preparation of antibody-coated coverslips

The glass coverslips were first coated with Polyornithine (10%) at room temperature for 1 hour. For antibody-coating, either LFA-1 antibody alone (9 μg/ml) or a combination of LFA-1 (9 μg/ml), CD3 (30 μg/ml) and CD28 (90 μg/ml) antibodies were applied. The antibodies were coated either at 30°C for 30 min or at 4°C overnight.

### Atomic force microscopy-based elasticity mapping in combination with light microscopy

Microscopic observation during elasticity mapping was carried out on a DMI 4000 B inverted microscope (Leica) with a 200-fold magnification. For elasticity mapping, primary T lymphocytes were suspended in AIM V without FCS, and seeded on antibody-functionalized coverslips at 37°C in presence of 5% CO_2_ for 15 min prior to elasticity mapping using a Bioscope Catalyst (Bruker) in Peak Force Quantitative Nanoscale Mechanical Characterization mode (Peak Force QNM) (Berquand et al., 2010). For the inhibitor treatment, T cells were pre-treated with the indicated inhibitor or vehicle control (DMSO) at room temperature for 30 min. Prior to each experiment, the AFM cantilever (MLCT cantilever B, Bruker) was calibrated using the thermal tune method (Li, Steinmetz et al., 2020). Elasticity mapping was carried out with the following parameters: line scan rate: 0.25 Hz, feedback gain: 0.5, peak force amplitude: 100 nm, peak force threshold: 700 pN and a resolution of ~60 nm. Young’s moduli were obtained by a Derjaguin-Muller-Toporov (DMT) fit (Derjaguin, Muller et al., 1975) of the retract part of each single force/distance curve. Elasticity maps (square-shaped with side length of 5-10 μm) spanning approximately a quarter of the cell, including lamellipodium and cell body, were recorded. Elastic moduli of the T-cells were determined as square shaped surface segments located on the cell bodies and lamellipodia. For lamellipodia, three individual square-shaped surface segments of 500 ×500 nm were analyzed per cell. If very slender filopodia structures with a lateral width of less than 500 nm were seen, the analyzed segment size was reduced to 250 × 250 nm. To determine the elastic moduli of the cell bodies, one 1.5 x 1.5 μm surface segment of the peripheral region was investigated per cell. Approximately 29,600 individual elasticity values were analyzed on a total of 58 primary T lymphocytes and Jurkat T-cells, respectively.

### Statistical analysis

Data are presented as mean ± SEM. GraphPad Prism 6 Software (San Diego, CA, USA) was used for statistical analysis. The differences between two groups were analyzed by Wilcoxon matched-pairs signed rank test. P values < 0.05 were considered significantly different. Individual *p* values are given in the figures.

## Author contributions

P.J. performed AFM-light microscopy measurements and determined the elasticity; X.Z. prepared the cells and antibody-coated coverslips and performed statistical analysis between groups; M.B. helped to analyze and interpret AFM data; B.Q. generated concepts, designed experiments, and wrote the manuscript; all authors contributed to the writing of the manuscript and provided advice.

## Acknowledgments

We thank the Institute for Clinical Hemostaseology and Transfusion Medicine for providing donor blood; Carmen Hässig, Cora Hoxha, Gertrud Schwär and Sandra Janku for excellent technical help. This project was funded by the Deutsche Forschungsgemeinschaft (SFB 1027 Project A2 to B.Q. and B2 to M.B.), INM Fellow (to B.Q.). The authors declare no competing interests.

## Ethical considerations

Research carried out for this study with healthy donor material (leukocyte reduction system chambers from human blood donors) is authorized by the local ethic committee (declaration from 16.4.2015 (84/15; Prof. Dr. Rettig-Stürmer)).

## Notes

### Competing Interest Statement

The authors have declared no competing interest.

